# Healthy B cells: allies or adversaries of CAR-T cell immunotherapy?

**DOI:** 10.1101/2024.11.08.622644

**Authors:** Emanuelle A. Paixão, Luciana R. C. Barros, Artur C. Fassoni, Regina C. Almeida

## Abstract

Chimeric antigen receptor (CAR)-T cell immunotherapy has achieved significant success against various haematological cancers, including B-cell malignancies. Its efficacy against B-cell cancers is influenced by the presence of healthy B-cells expressing the target antigen, and B-cell aplasia (BCA) serves as an indicator of successful therapy outcome. However, the precise influence of healthy B-cells on the *in vivo* dynamics of CAR-T cells and their ultimate impact on therapy outcomes remain unclear. Here, we propose a mathematical model to describe CAR-T cell immunotherapy in B-cell cancer patients. Our model successfully captured the interactions between different CAR-T cell phenotypes, tumour cells, and healthy B cells in patients who achieved a complete response. Using these cases, we constructed virtual scenarios to investigate how variations in baseline tumour and healthy B-cell populations, along with patient-specific factors related to CAR-T cell expansion and B-cell influx from the bone marrow, affect treatment outcomes. Our results suggest that the onset and duration of BCA is a patient-specific feature that depends primarily on the continuous influx of newly generated B cells, their proliferative capacity, and the expansion and cytotoxicity of CAR-T cells.

**Statement of significance:** This study presents a significant advancement in understanding the dynamics of CAR-T cell immunotherapy in B-cell malignancies by introducing a mathematical model that captures the complex interactions between CAR-T cells, tumour cells, and healthy B cells. The model provides crucial insights into how patient-specific factors, such as baseline tumour burden, B-cell populations, and CAR-T cell expansion, influence treatment outcomes, including the onset and duration of B-cell aplasia (BCA), a key marker of therapeutic success. Healthy B cells can act as allies, adversaries, or have a neutral effect on the therapy, depending on the tumour burden. These findings highlight the importance of personalised approaches in CAR-T cell immunotherapy, offering potential pathways to optimise treatment strategies for improved efficacy and patient outcomes.

## Introduction

CAR-T cell immunotherapy is an adoptive therapy that has achieved significant success in treating haematological cancers; however, high recurrence rates continue to pose a substantial challenge (1, 2). A critical factor in its success is the selection of an appropriate target antigen expressed by the tumour cells. Ideally, the chosen antigen should exhibit high expression levels on tumour cells while being absent or minimally expressed on normal, especially vital, cells. In the context of B-cell cancers, various B-cell lineage-specific antigens, such as CD19, CD20, CD21, CD22, CD23, CD72, and surface immunoglobulin, are potential targets. Among these, the CD19 antigen is particularly suitable for CAR-T cell immunotherapy due to its consistently high expression across B-cell neoplasms and its absence on non-B-cell lineage cells, including haematopoietic stem cells, plasma cells, and other healthy tissues. Moreover, although CD19-targeted CAR-T cell therapy affects both malignant and normal CD19-positive cells, healthy B cells can be regenerated through stem cell differentiation in the bone marrow, eventually restoring their homeostatic levels post-therapy (3, 4).

Unlike immunodeficient mice, applying CAR-T cell immunotherapy in patients involves complex interactions with the host immune system and non-tumour cells. These interactions can significantly impact the therapeutic dynamics in B-cell malignancies. Importantly, normal B lymphocytes express the target antigen, contributing to the *in vivo* expansion of CAR-T cells and modulating their cytotoxic activity. Evidence indicates a positive correlation between therapy responses and the total CD19-positive antigen load, including both tumour and normal B-cells, in patients prior to therapy (5). Consequently, B-cell aplasia (BCA) serves as an indicator of the persistence of functional CAR-T cells and successful therapeutic outcomes. Monitoring BCA can guide clinical observations on disease progression and inform decisions about further interventions, such as allogeneic haematopoietic stem cell transplants (6).

Gardner et al. (5), in their study of children and young adults with relapsed/refractory B-cell acute lymphoblastic leukaemia (R/R B-ALL) treated with CAR-T19 BB*ζ*, found a positive correlation between the magnitude and peak engraftment of CAR-T cells in the peripheral blood and total CD19 load in the bone marrow, but no correlation with infused cell dose or disease burden at infusion. They observed CAR-T cell peaks around day 10, with a range spanning from 7 to 18 days, remission within 21 days, and a median BCA duration of 3 months. In a study with CAR-T19 28*ζ*, Lee et al. (7) reported strong anti-leukaemic activity in children and young adults with chemotherapy-resistant B-ALL, with CAR-T cell expansion peaking between 7 and 14 days and B-cell recovery by day 28 in some patients. Higher CAR-T cell expansion was linked to a complete response, and the cohort had a median overall survival of 10.5 months (8). Therefore, successful CAR-T cell therapy typically shows circulating malignant blasts at therapy onset, CAR-T cell expansion peak around 1-2 weeks with concurrent elimination of tumour and healthy B cells, undetectable CAR-T cells by day 50, and recovery of normal B cells with sustained absence of malignant blasts (7). Wang et al. (9) conducted a long-term follow-up study with 115 children and young adults with R/R B-ALL treated with CD19 CAR-T cells. Of these, 112 cases exhibited BCA with a median BCA persistence time of 30 days (ranging from 11 to 933 days) after infusion. In patients who did not undergo transplantation, the predictive value of BCA for outcomes strengthened over time. However, this indicator has limitations, as some patients experienced CD19-negative relapse.

Although CAR-T cell therapy has opened an avenue for mathematical modeling, very few studies have focused on considering the dynamic interactions with healthy B cells in patients (10–12). In Martínez-Rubio et al. (11), a model consisting of seven ordinary differential equations (ODEs) was proposed to describe the dynamics of leukaemic cells, effector and memory CAR-T cells, and four subpopulations of healthy B lymphocytes. These B cells comprised different maturation stages, three of which expressed the target antigen. The authors concluded that the dynamics of CAR-T cells and the response to therapy are independent of the infused dose or the initial burden of both target-antigen-expressing tumour and normal cells. The study by Serrano et al. (12) extended the work of (10), modelling the interaction between CAR-T cells, tumour cells, and healthy B cells. Focusing on stability and bifurcation analysis, this research demonstrated that a steady influx of healthy B cells is crucial for tumour eradication.

Here, we developed a model to describe CAR-T cell immunotherapy in patients with B-cell cancers, considering the interplay between different CAR-T cell phenotypes with target-antigen-expressing tumour and healthy B cells. Focus is given on the analysis of target-antigen-dependent mechanisms and their role in the response to therapy.

## Methods

### Mathematical model

We have recently developed a model that describes the dynamic interaction between various CAR-T cell phenotypes and tumour cells (13). This model successfully captured the multiphasic dynamics observed following CAR-T cell infusion in patients with different haematological cancers and therapeutic outcomes. Here, we extend this existing model by incorporating the interactions of CAR-T and tumour cells with target-antigen-expressing healthy B cells. The main assumptions of the proposed model are:

(H1) CAR-T cells undergo clonal expansion when in contact with target-antigen-expressing tumour and healthy B cells (14, 15).
(H2) The conversion (or reactivation) of memory to effector CAR-T cells depends on the contact between memory CAR-T cells with target-antigen-expressing tumour and healthy B cells (16).
(H3) Tumour and healthy B cells compete for shared resources, with tumour cells having a stronger competitive advantage (17).
(H4) Tumour cells exhibit increased proliferation compared to healthy B cells, due to the acquisition of functional capabilities developed during their evolution (18).
(H5) A constant flow of newly generated healthy B cells continuously enters the peripheral blood from the bone marrow (19).
(H6) The cytotoxic effects of CAR-T cells on tumour and healthy B cells differ, with the impact on healthy B cells surpassing that on tumour cells. This discrepancy arises from immune evasion and resistance mechanisms employed by tumour cells (20, 21), such as temporarily hiding target antigens (22, 23).

An overview of the model interaction mechanisms is provided in Figure 1, with the model equations detailed in Box 1. A brief description of the model assumptions is given below; further details can be found in (13).

**Figure 1.**
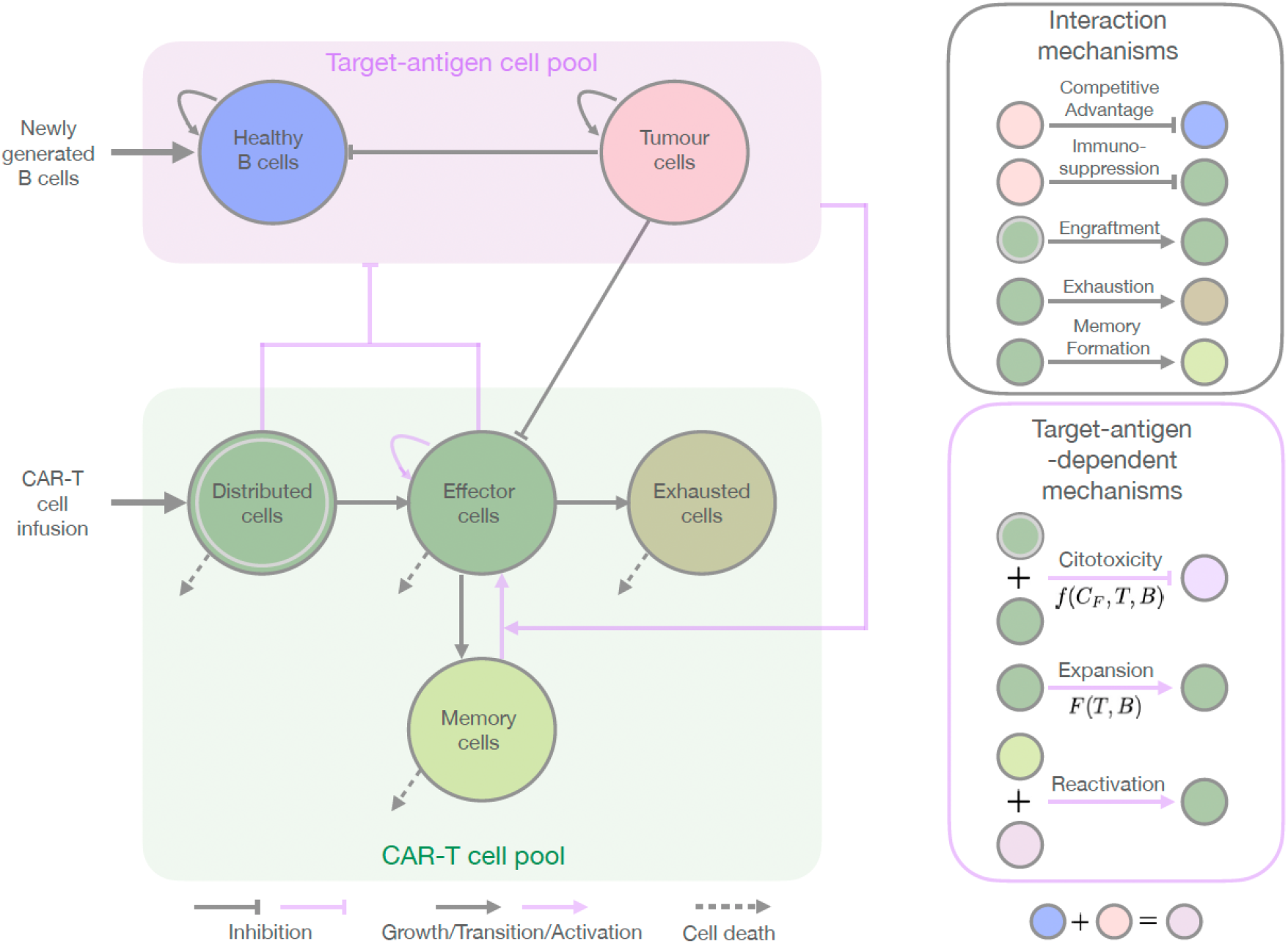
Schematic description of the CAR-T cell immunotherapy model in patients (left) and the corresponding interaction and target-antigen-dependent mechanisms (right). Interactions influenced by the target antigen are denoted by light magenta lines. The CAR-T cells, specifically designed to target the chosen antigen, undergo a rapid distribution phase upon injection into the patient. During this phase, they spread throughout the body and die naturally (*C*_*D*_). A subset of these cells undergoes engraftment and successfully integrates into the bloodstream and tumour niche. Referred to as effector CAR-T cells (*C*_*T*_), they expand upon target antigen contact, exhibit cytotoxicity against cells expressing the target antigen (both tumour and healthy B cells), differentiate into memory CAR-T cells, and are impaired by tumour-induced immunosuppressive mechanisms. They may also die naturally or become exhausted, losing functionality. Both *C*_*T*_ and *C*_*D*_ populations exhibit cytotoxic effects and are consequently designated as functional CAR-T cells (*C*_*F*_). The long-term memory CAR-T cells (*C*_*M*_) may naturally undergo death or remain highly responsive to cells expressing the target antigen. Upon interaction with tumour or healthy B cells, they undergo reactivation into effector CAR-T cells, producing a rapid immune response against the tumour. Over time, effector CAR-T cells become exhausted (*C*_*E*_) and undergo apoptosis. Tumour (*T*) and healthy B (*B*) cells grow subject to intra- and interspecific competitions while sharing the same carrying capacity. Owing to a stronger competitive advantage, tumour cells inhibit the growth of healthy B cells. Nonetheless, a continuous influx of newly generated B cells from the marrow facilitates the eventual repopulation of healthy B cells.

The populations of tumour and healthy B cells are denoted by *T* and *B*, respectively. The CAR-T cell population includes four phenotypes, referred to as distributed (*C*_*D*_), effector (*C*_*E*_), memory (*C*_*M*_), and exhausted (*C*_*E*_) cells. After the infusion of a dose of *C*_*D*_(0) CAR-T cells, these distributed CAR-T cells die at a rate of *β* and transition to the effector phenotype with an engraftment rate of *η*. The total number of engrafted CAR-T cells (*EC*) is approximately given by

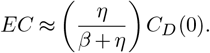

The condition *EC* ≥ 1 must hold to ensure that at least one cell undergoes engraftment (13). In some cases, only a fraction of the injected CAR-T cells may engraft, survive, expand, and generate memory (24, 25). Although our model does not explicitly consider the number of clones, we model the population of effector CAR-T cells as deriving from the engraftment of a variable number of CAR-T cells.

Once engrafted, effector CAR-T cells expand at a time- and antigen-dependent rate of *κ*(*t*)*F* (*T, B*). The time-dependent rate *κ*(*t*) is defined as:

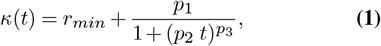

where the parameters *r*_*min*_, *p*_1_, *p*_2_, and *p*_3_ describe patient and product characteristics related to the basal expansion rate, strength, duration, and decay of CAR-T cell expansion, respectively (13). The function *F* (*T, B*) describes the regulation of CAR-T cell expansion by target-antigen-expressing cells (H1) and is defined as

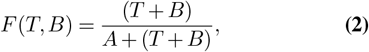

in which *A* is a half-saturation constant.

Target-antigen-expressing tumour and healthy cells also drive the conversion of memory to effector phenotype at a rate of *θ* (H2). In addition, effector CAR-T cells decline due to immunosuppressive mechanisms induced by tumour cells at rate of *α*, as well as due to natural death and differentiation to memory and exhausted phenotypes at rates *ξ, ϵ*, and *λ*, respectively. Exhausted and memory CAR-T cells have death rates of *λ* and *µ*, respectively.

Cells expressing the target antigen, including both tumour and healthy B cells, experience competitive effects for available resources, characterised by the carrying capacity (1*/b*) (further details may be found in the supplementary material – SM). However, given the stronger competitive advantage of tumour cells over healthy B cells, we assume that the competition between these cells adversely affects only the population of healthy B cells, modulated by the coefficient *ω* (H3). Tumour cells exhibit a greater proliferative capacity compared to healthy B cells, establishing a constraint between their maximum growth rates in the form *r*_1_ *> r*_2_ (H4). A constant influx of B-cell precursors from the marrow is represented by *B*_*p*_ (H5).

Tumour and healthy B cells are killed by CAR-T cells at rates of *γ*_1_*f* (*C*_*F*_, *T, B*) and *γ*_2_*f* (*C*_*F*_, *T, B*), respectively. The cytotoxic effect of functional CAR-T cells on healthy B cells surpasses that on tumour cells, implying that *γ*_1_ *< γ*_2_ (H6). Here, *C*_*F*_ = *C*_*D*_ + *C*_*T*_ represents the number of functional CAR-T cells that are activated to kill target-antigen-expressing cells. The functional response *f* (*C*_*F*_, *T, B*) is given by

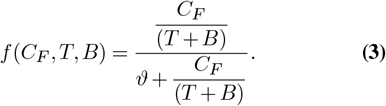

This function characterises the killing process as dependent on the binding of functional CAR-T cells to target-antigen-expressing cells, with a half-saturation constant *ϑ* (13, 26).

The biological meanings and units of the parameters are shown in Table 1.

**Table 1.** Model parameters with their units and biological meanings, as well as constraints applied to specific parameters.

| Parameter | Unit | Biological meaning |
| --- | --- | --- |
| $\beta$ | day<sup>-1</sup> | Reduction rate of infused cells due to natural death during their distribution in the patient's body |
| $\eta$ | day<sup>-1</sup> | Engraftment rate of injected cells to blood and tumour niche |
| $r_{min}$ | day<sup>-1</sup> | Minimum expansion rate of effector CAR-T cells |
| $p_1$ | day<sup>-1</sup> | Initial expansion rate of effector CAR-T cells |
| $p_2$ | day<sup>-1</sup> | Rate that regulate the duration of maximum expansion period of effector CAR-T cells |
| $p_3$ | - | Expansion coefficient that regulates the decay of maximum expansion period of effector CAR-T cells |
| $A$ | cell | Half-saturation constant of function $F(T, B)$ |
| $\xi$ | day<sup>-1</sup> | Death rate of effector CAR-T cells |
| $\epsilon$ | day<sup>-1</sup> | Differentiation rate of effector CAR-T cells into memory CAR-T cells |
| $\lambda$ | day<sup>-1</sup> | Exhaustion rate of effector CAR-T cells |
| $\theta$ | (cell.day)<sup>-1</sup> | Reactivation coefficient of memory CAR-T cells into effector CAR-T cells due to interaction with target-antigen-expressing cells |
| $\alpha$ | (cell.day)<sup>-1</sup> | Inhibition coefficient of effector CAR-T cells due to interaction with tumour cells |
| $\mu$ | day<sup>-1</sup> | Death rate of memory CAR-T cells |
| $\delta$ | day<sup>-1</sup> | Death rate of exhausted CAR-T cells |
| $r_1$ | day<sup>-1</sup> | Maximum growth rate of tumour cells |
| $\gamma_1$ | day<sup>-1</sup> | Cytotoxic rate of the functional CAR-T cells on tumour cells |
| $\vartheta$ | - | Half-saturation constant of the cytotoxic effect on antigen expressing cells |
| $r_2$ | day<sup>-1</sup> | Maximum growth rate of healthy B cells |
| $b$ | cell<sup>-1</sup> | Inverse of the tumour and healthy B cell carrying capacity |
| $B_p$ | cell.day<sup>-1</sup> | Total B cell precursors, expressing target antigen, generated in bone marrow |
| $\gamma_2$ | day<sup>-1</sup> | Cytotoxic rate of the functional CAR-T cells on healthy B cells |
| $\omega$ | (cell.day)<sup>-1</sup> | Competition coefficient between tumour and healthy B cells |
| Constraint | Meaning |  |
| $\xi > \mu$ | Memory CAR-T cells have a longer lifetime than effector CAR-T cells | |
| $\left(\frac{\eta}{\beta + \eta}\right) C_D(0) \geq 1$ | The total number of engrafted CAR-T cells ($EC$) ranges from one cell to the total dose of CAR-T cells | |
| $r_1 > r_2$ | Tumour cells exhibit a higher proliferative capacity compared to healthy B cells | |
| $\gamma_1 < \gamma_2$ | The cytotoxic effects of functional CAR-T cells on healthy B cells are greater than on tumour cells. This is primarily due to the mechanisms that tumour cells use to evade immune system surveillance | |

#### Box 1

**System of six ordinary differential equations**

Distributed CAR-T cells 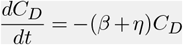

Effector CAR-T cells 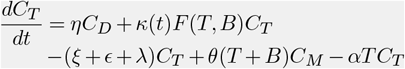

Memory CAR-T cells 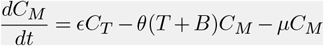

Exhausted CAR-T cells 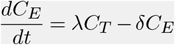

Tumour cells 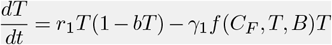

Healthy B cells 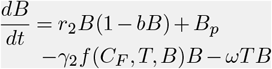

### Experimental data and calibration

To assess the model’s capability to describe the impact of healthy B cells on therapy dynamics, we surveyed the literature for datasets presenting the temporal dynamics of CAR-T cells alongside tumour and healthy B cells. We identified a single study by Lee et al. (7) that provided median data for these three cell populations. Subsequently, we searched for other works that described at least the dynamics of CAR-T cells and healthy B cells. In this context, we selected the work published by Kochenderfer et al. (27) for providing data from both populations in the same measurement unit (cells/*µ*L). The chosen datasets were extracted using the WebPlotDigitizer software (28) and are provided in the Tables SM.1–3.

The first dataset (7) pertains to paediatric and young adult patients (ages 1–30 years) with R/R B-ALL or non-Hodgkin lymphoma. It comprises median data from the group of responding patients, measured in absolute circulating cells. All included patients underwent chemotherapy prior to receiving a single dose of autologous second-generation anti-CD19 CAR-T cell constructs with a CD28 co-stimulatory domain. Doses were reported in cells/kg and converted to cell counts, assuming an average patient weight of 60 kg. The clinical limit of detection for all cell populations is 10^4^ cells.

The second dataset (27) includes data from four adult patients with different subtypes of diffuse large B-cell lymphoma (DLBCL). These patients, referred to as *Ki*, with *i* corresponding to the same patient numbering used in (27), underwent chemotherapy followed by the administration of autologous anti-CD19 CAR-T cells with a CD28 costimulatory domain. Individual doses were also reported in cells/kg, and an average patient weight of 60 kg was assumed for dose calculations. All patients received a single dose of CAR-T cells and achieved long-term complete response of varying durations, ranging from 38 to 56 months. The *in vivo* CAR-T and healthy B cell data were reported in peripheral blood mononuclear cells (PBMCs) per *µ*L, which were converted to cell counts using the transformation reported in (13, 29) (1 cell/*µ*L corresponds to 5×10^8^ cells). The detection threshold for healthy B cells was indicated as 60 cells/*µ*L. The detection limit for CAR-T cells varies for each patient, as it was given by the percentage of PBMCs containing the CAR gene. The PBMC counts are assumed here to represent the peak CAR-T cell values. The resulting detection thresholds are likewise applied to tumour cell counts.

Examining the underlying mechanisms that regulate each phase of CAR-T cell activity, a systematic approach for calibrating patient-specific models using experimental data was established in (13). The current study employs the same methodology to define an initial parametric space. The first step involved fine-tuning the parameters to capture the dynamics of CAR-T cells. Subsequently, we refined the existing parameter set to incorporate the dynamics of healthy B cells. This process was repeated to account for the collective dynamics of all cell populations. During this step, given that a complete response was observed in all cases, the search for the parameters was carried out by ensuring that the tumour burden remained below the detection threshold on the final day of follow-up. Through this procedure, after conducting exhaustive simulations, we successfully captured the scenarios outlined in the studies by Lee et al. (7) and Kochenderfer et al. (27) using the (non-unique) parameter sets presented in Table SM.4.

### Model settings and numerical solution

The explicit fourth-order Runge–Kutta method (30) was employed for the numerical solution of the model outlined in Box 1. The code was implemented in the C programming language. After conducting multiple tests to determine the most suitable and convergent time step size, we set Δ*t* = 10^−6^ day.

For the initial conditions, the CAR-T cell dose was assigned as the starting value for the distributed CAR-T cell population (*C*_*D*_(0)), with null values for the other CAR-T cell phenotypes (*C*_*T*_ (0) = *C*_*M*_ (0) = *C*_*E*_(0) = 0 cell). In cases where baseline conditions for the tumour and/or healthy B cell populations were unavailable, we assumed *T* (0) = 10^7^ cells and/or *B*(0) = 10^7^ cells.

Throughout the simulations, therapy responses are assessed by examining the tumour burden on the final day of analysis according to the following criteria: for complete response (CR), the tumour burden must fall below the detection threshold; for partial response (PR), the tumour burden should be above the detection threshold but below 50% of the initial tumour burden; for stable disease (SD), the tumour burden must remain within ± 50% of the initial tumour burden; and for progressive disease (PD), the tumour burden must exceed ±50% of the initial tumour burden (31, 32).

## Results

The success of CAR-T cell immunotherapy is influenced by specific characteristics of the CAR-T cell product, primarily its expansion capacity within the body. Here, applying a mathematical model for CAR-T cell therapy encompassing the interactions of tumour and healthy B cells, we suggest that the initial composition of the total target-antigen load is also a crucial factor. Variations in the initial densities of tumour and healthy B cells lead to different outcomes as this composition affects biological mechanisms such as cytotoxicity, competition, and immunosuppression.

### Modelling different scenarios with complete response to CAR-T cell immunotherapy

First, we analysed the model dynamics by fitting it to the median patient data from (7), and evaluated the model’s accuracy in capturing the time courses of CAR-T, tumour, and healthy B cells (Figure 2, left column – *Profile CR90*). The CAR-T cell population exhibits typical multiphasic dynamics. Following the injection of an estimated dose of 6×10^7^ CAR-T cells, we observed the *Distribution* phase, characterised by a rapid decline in CAR-T cells alongside their immediate cytotoxic effect on tumour and healthy B cells. This is followed by the *Expansion* phase, during which the engrafted CAR-T cells (approximately 2.57×10^3^ cells) proliferate, leading to a more pronounced reduction in the target-antigen-expressing cells. This expansion is modulated by the function *κ*(*t*). Until approximately the sixth day after infusion, the maximum expansion rate of 2.01 day^−1^ is maintained, gradually declining to a minimum value of 0.10 day^−1^ around a month after infusion (Figure 2, left column, bottom row). CAR-T cells peak on day 9 with 2.89×10^7^ cells, predominantly in the effector phenotype. Significant exhaustion of these cells is observed, but there is also the generation of a clinically detectable pool of memory cells. The subsequent *Contraction* phase is characterised by a rapid decline in circulating CAR-T cells due to the programmed cell death of activated CAR-T cells and the loss of antigenic stimulation. This decline continues at a slower pace beyond 45 days, sustained by the presence of memory cells.

**Figure 2.**
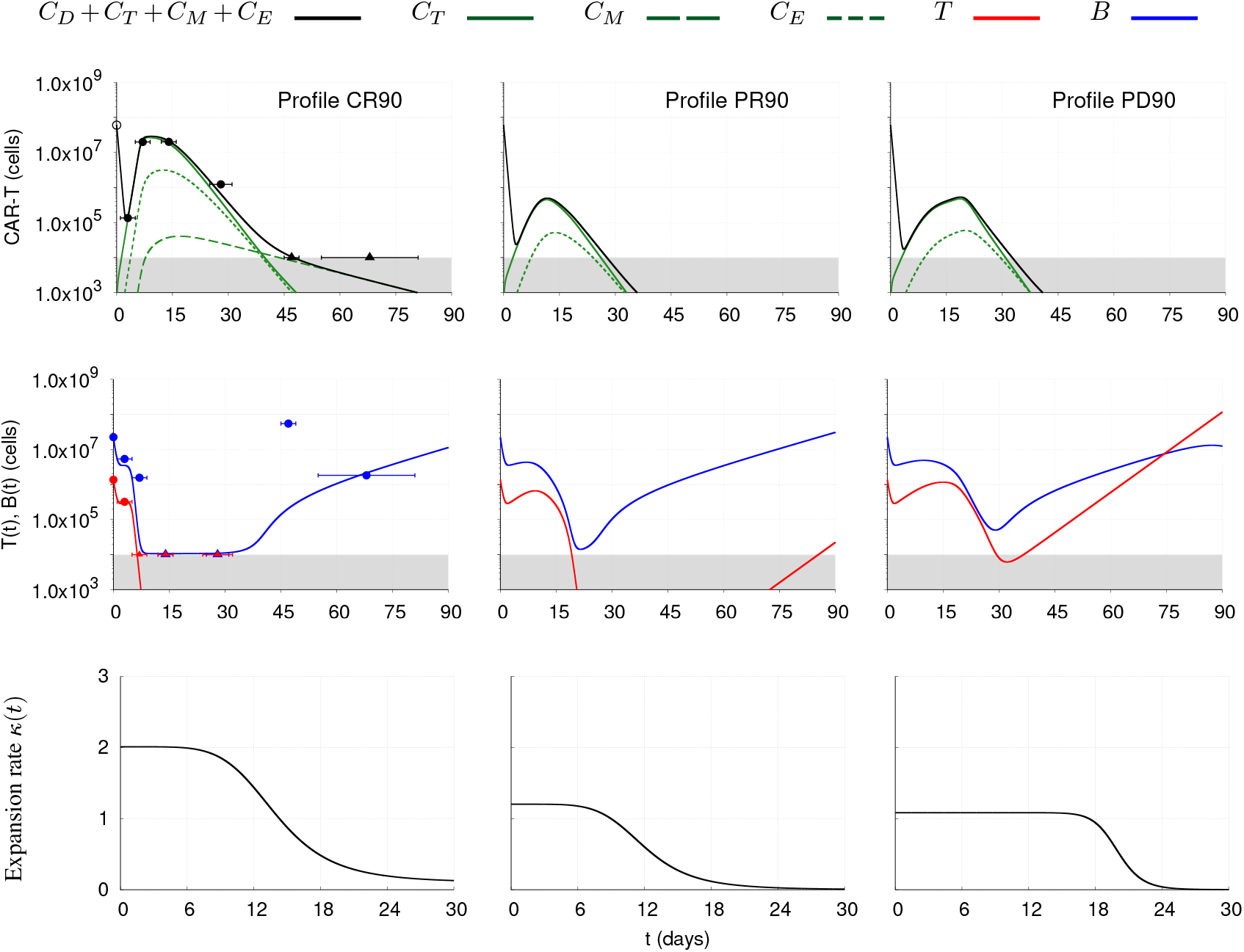
Model simulations for the median responding patient reported in Lee et al. (7) (left column), who achieved complete response (CR) at the last follow-up (interval from infusion to the last follow-up in days). Experimental data are shown as circular dots for values above the detection threshold of 10^4^ cells and as triangles for values below this threshold. Error bars indicate variations (range) in the days of the measurements: days 3 (1–5), 7 (5–9), 14 (12–16), 28 (25–31), 42 (45–49), and 68 (55–81). The median dose value of 6.0*×*10^7^ cells (∘) was estimated based on patients with an average weight of 60 kg. Theoretical patient profiles were generated by modifying the *κ*(*t*) function, resulting in different therapeutic results 90 days after therapy: Profile PR90 (median column), indicating partial response, and Profile PD90 (right column), indicating progressive disease. The total CAR-T cell population 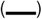 is divided into effector (*C*_*T*_), memory (*C*_*M*_), and exhausted (*C*_*E*_) phenotypes, represented by continuous, dashed, and dotted green lines, respectively. Dynamics of tumour 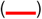 and healthy B cells 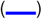 are also presented. The gray region represents undetectable levels (below the threshold of 10^4^ cells). The corresponding time-dependent expansion rate function (*κ*(*t*)) is shown for each profile.

In analysing the dynamics of target-antigen-expressing cells, we observed a significant impact from the continuous supply of healthy B cells from the bone marrow (*B*_*p*_). By day 7, tumour cells became undetectable, showing a sharp decline, while healthy B cells remained detectable, albeit at a density near the clinical detection threshold. In the model simulation, B-cell aplasia (BCA) was not observed, as the healthy B-cell population reached a minimum of 1.06×10^4^ cells—close to, but not below, the detection threshold. While functional CAR-T cells remained at a high density, *B* cells were controlled; however, as CAR-T cells declined, healthy B cells began to regrow due to the constant influx of *B*_*p*_. In contrast, tumour cell numbers stayed low and clinically undetectable until day 90, indicating a complete response. When extending the simulation over a longer period (not shown), the healthy B cell population returned to its homeostatic level, nearing carrying capacity, approximately one year after infusion. A highly discrepant data point in the healthy B cell population appears on day 47, likely an outlier due to the reduced number of evaluated patients, as indicated by the experimental data in Table SM.1.

The proposed model was also fitted to patient-specific data from Kochenderfer et al. (27) (Figures SM.1–4), including four patients with different subtypes of DLBCL and distinct temporal dynamics. DLBCL is a type of cancer characterised by a highly pronounced CAR-T cell distribution phase (33). Despite the lack of experimental data within this phase, our model successfully captured the CAR-T cell dynamics of the four patients analysed, showing distinct distribution phases with varying intensities and durations as well as different BCA durations. Detail analyses are provided in the SM.

### Heterogeneities in patients and CAR-T cell products induce different outcomes

After demonstrating the model’s ability to fit both median and patient-specific data, we aimed to evaluate the impact of both patient and infused product heterogeneities on response dynamics. Based on the *Profile CR90* of the median responding patient, we generated two theoretical patients, denoted as *Profile PR90* and *Profile PD90* (Figure 2 – middle and right columns), by modifying their functions *κ*(*t*), while keeping constant all other parameters. Both theoretical profiles were designed with a lower maximum expansion rate than *Profile CR90*, with *Profile PR90* maintaining a similar duration for the expansion plateau (∼ 6 days), while *Profile PD90* exhibited a longer duration (∼ 18 days) (Figure 2 - bottom row). For these designed theoretical patients, the slight variations in CAR-T cell expansion profiles resulted in different dynamics of the target-antigen-expressing cells, ultimately leading to different responses to therapy. Indeed, the slightly higher maximum expansion rate in *Profile PR90* resulted in an undetectable number of tumour cells after the expansion phase; however, this did not prevent tumour cells from reappearing at clinically detectable levels at day 90, indicating a partial response (PR90). In contrast, the prolonged but lower expansion profile of the function *κ*(*t*) in *Profile PD90* allowed for greater persistence of tumour cells, and by day 90, the response to therapy is classified as progressive disease (PD90).

These varying responses to therapy, resulting solely from differences in the function *κ*(*t*), suggest that heterogeneities in the infused products and patient-specific factors influencing CAR-T cell dynamics — particularly the intensity and duration of expansion — have a substantial effect on the overall therapeutic response.

### Effect of initial target-antigen load composition on therapy outcomes

It is known that the total target-antigen load has a direct impact on the expansion and cytotoxicity of CAR-T cells, thereby influencing therapeutic outcomes (34, 35). In this context, it is important to understand how these mechanisms are affected by the composition of the total target-antigen load, specifically the fractions consisting of tumour and healthy B cells. To shed light on this question, we evaluated several combinations of initial conditions *T* (0) and *B*(0) within the range of 10^4^ to 10^9^ cells, as determined by the experimental data from Lee et al. (7). We then applied these different initial conditions to the three previously described profiles (*CR90, PR90*, and *PD90*), while keeping all other parameters constant. In this analysis, we also included the model fit for patient 7 from (27), denoted as *Patient K7*, whose dynamics are presented in Figure SM.3. For each simulation, we recorded the theoretical (predicted) relapse-free time (*t*_*T R*_), defined as the time until the tumour relapses after achieving complete remission. The results of these *in silico* experiments are presented in Figure 3.

**Figure 3.**
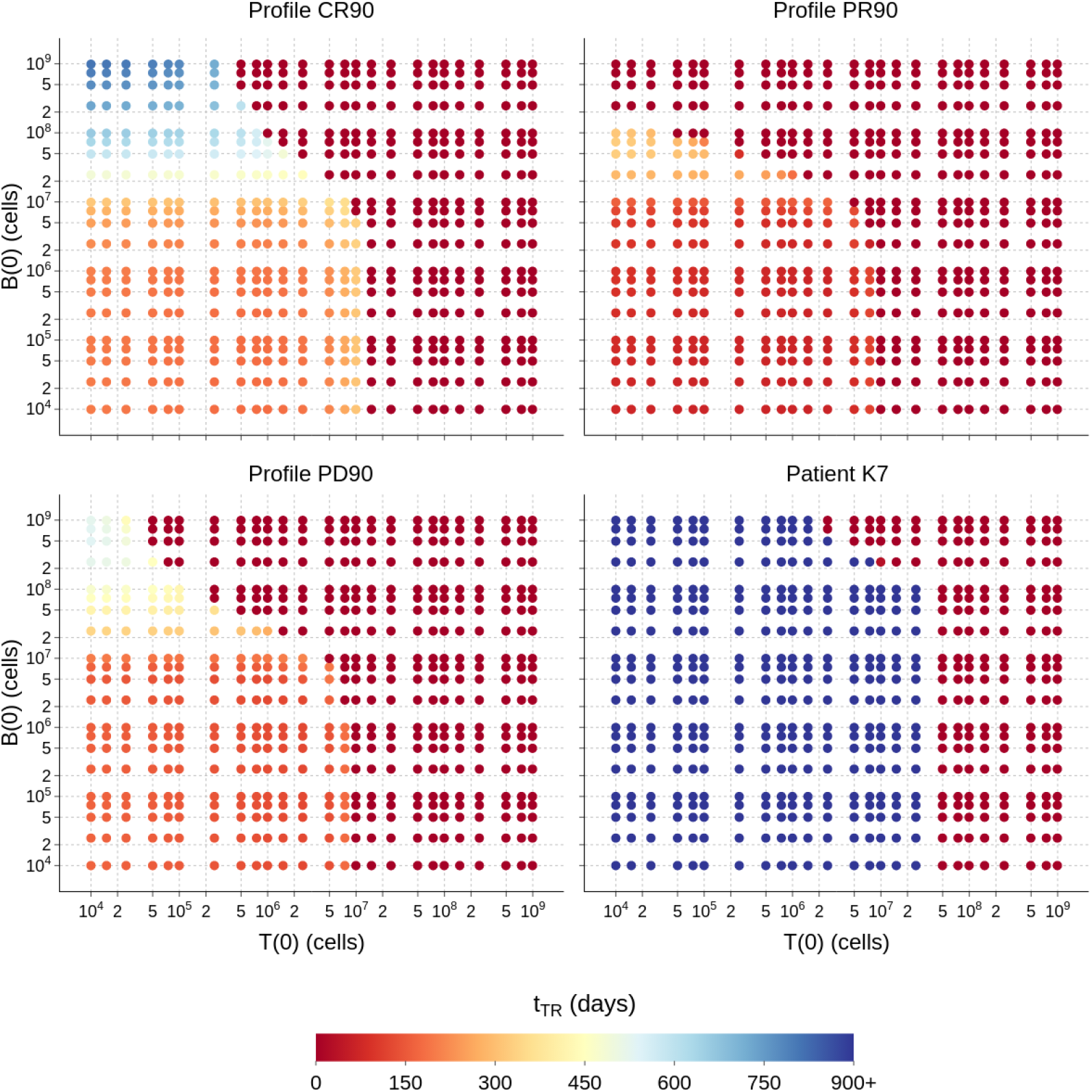
Theoretical relapse-free time (*t*_*TR*_) recorded for profiles CR90, PR90, PD90, and patient K7. For each scenario, only the initial conditions were varied in the range of 10^4^ to 10^9^, while maintaining the values of the model parameters. A null value indicates that the tumour cell count did not fall below the clinical detection threshold. On the other hand, (*t*_*TR*_ = 900+) represents recurrence within 900 days or more.

The results for *Profile CR90* revealed three distinct regions (yellowish, blueish, dark red). At low tumour burden, increasing *B*(0) led to later relapse-free times, indicating that healthy B cells contribute positively to the therapy, acting as important allies. As the initial tumor burden increased, this beneficial effect of healthy B cells became less pronounced. For *T* (0) between 5×10^5^ and 1×10^7^ cells, we observed a *B*(0) value beyond which *t*_*TR*_ = 0 days, indicating treatment failure, as the tumour cell population no longer decreased below the clinical detection threshold. At very high tumor burdens exceeding 1.5×10^7^ cells, treatment ultimately failed, and the initial number of healthy B cells had no impact on the therapy outcome. Moreover, for healthy B cell counts below 5×10^6^ cells, the relapse-free time did not vary linearly with an increase in tumour burden. Interestingly, there was an improvement in therapy response when *T* (0) was near 10^7^ cells. However, just below this value, *t*_*TR*_ decreased, while therapy ultimately failed for *T* (0) ≥ 1.5×10^7^ cells. It is also noteworthy that an equivalent total initial target-antigen load, even with different compositions of *T* (0) and *B*(0) cells, resulted in varying relapse-free times. For instance, with *B*(0) = 10^4^ cells and *T* (0) = 10^9^ cells, there was no response to therapy. In contrast, with *T* (0) = 10^4^ cells and *B*(0) = 10^9^ cells, a positive response was observed, yielding a relapse-free time of *t*_*TR*_ = 800 days. This suggests the existence of a therapeutic window, from the orange to the blue region, where B cells contribute positively to the therapy when the tumour burden is low.

Since *Profile PD90* demonstrated a poorer response to therapy than *Profile PR90*, as shown in Figure 2, one would expect *Profile PD90* to also yield worse results when varying the initial conditions. However, while *Profile PR90* exhibited earlier relapse-free times, *Profile PD90* revealed a blueish region indicating improved responses. These findings suggest that a high concentration of healthy B cells, in combination with a low tumour burden, positively impacts patients who experience prolonged CAR-T cell expansion.

The results for patient K7 showed a markedly different pattern compared to the previous cases. As a patient with CR51^+^ months, as indicated in (27), the changes in the proportions between *T* (0) and *B*(0) revealed either treatment failure (in dark red) or success (in dark blue), with relapse-free times exceeding 900 days. Regardless of the number of healthy B cells, treatment success was achieved when *T* (0) *<* 2 × 10^6^ cells. Above this value, treatment failure happened at high concentrations of healthy B cells, with *T* (0) ≥ 5×10^7^ cells resulting in relapse-free times of *t*_*TR*_ = 0 day.

We next detailed the temporal dynamics of CAR-T cells, with their distinct phenotypes, alongside tumour and healthy B cells, for five scenarios with different initial conditions selected from *Profile CR90* in Figure 3. The behaviour of function *f* (*C*_*F*_, *T, B*), which modulates the cytotoxic effect, reflects the impact of the total target-antigen load on the therapy response (Figure 4 – top panel). Functional CAR-T cells determine when the function *f* (*C*_*F*_, *T, B*) is non-zero. By design, its highest values are obtained when *C*_*F*_ */*(*T* + *B*) *>* 1, which correlates with BCA. In scenarios 1 and 2, the initial CAR-T cell dose of 6×10^7^ cells and the total target-antigen load are insufficient to promote CAR-T cell expansion and tumour elimination. Although in scenario 3 the expansion of effector CAR-T cells is less than that in scenario 2, the ratio *C*_*F*_ */*(*T* + *B*) *>* 1 was sufficient to sustain cytotoxicity and reduce the tumour below the detection threshold. The greater the expansion of CAR-T cells and the longer the duration in which *C*_*F*_ */*(*T* + *B*) *>* 1, the longer the cytotoxic effect remains at its peak, and the more prolonged the period of BCA, as observed in scenarios 4 and 5.

**Figure 4.**
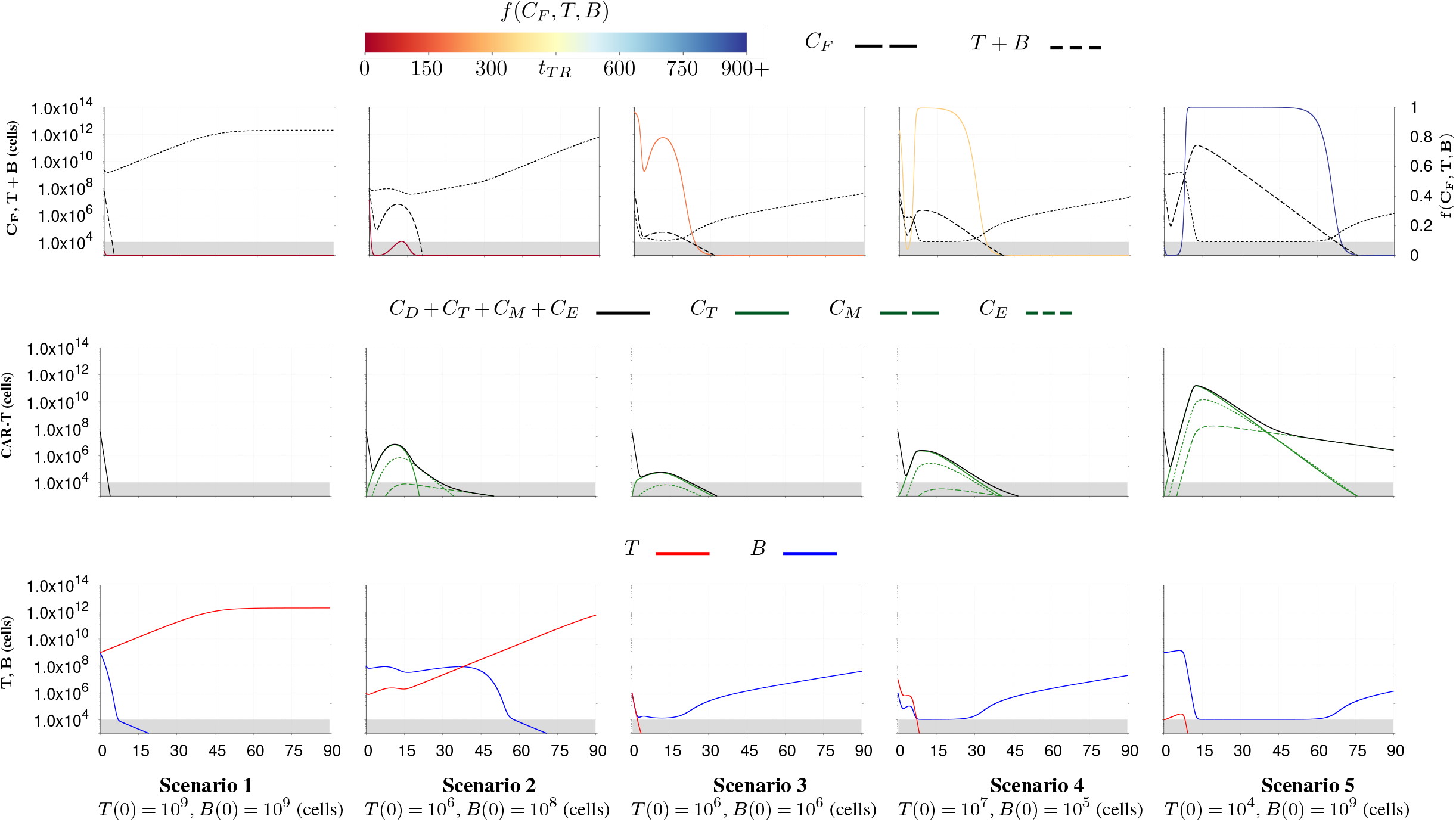
Model simulations for *Profile CR90*, using specific combinations of initial conditions for *T* (0) and *B*(0), with a dose of 6 *×* 10^7^ CAR-T cells. The behaviour of the function *f* (*C*_*F*_, *T, B*) in relation to distinct theoretical relapse-free time (*t*_*TR*_) is presented, alongside the total antigen load and the functional CAR-T cell population (top panel). The middle and bottom panels illustrate the temporal dynamics of CAR-T cell phenotypes, as well as tumour and healthy B cells, respectively.

Figure 4 shows that a successful therapeutic response is characterised by a rapid decrease in tumour burden (scenarios 3, 4, and 5). Otherwise, competition between populations expressing the target antigen becomes evident, resulting in a reduction in the healthy B cell population, as observed in scenarios 1 and 2. In scenario 5, a large memory pool was formed alongside significant expansion of effector CAR-T cells. This appears to favour the ratio between *C*_*F*_ and *T* +*B*, sustaining the cytotoxic effect and BCA for longer periods.

Our simulations indicate that the initial antigen load and its distribution between tumour and healthy B cells are crucial factors for successful therapy. However, variations in individual patient profiles and CAR-T cell product characteristics also significantly influence treatment outcomes.

### Interplay between initial target-antigen load composition and biological mechanisms

In addition to the cytotoxicity and expansion of CAR-T cells, all biological mechanisms modulated by the target-antigen load, such as immunosuppression and competition between tumour and healthy B cells, influence the therapy response to some degree. To analyse the impact of the initial composition of cells expressing the target antigen on these mechanisms, we examined the set of simulations presented in Figure 3 for *Profile CR90*. For these simulations, Figure 5 illustrates the dynamics of total CAR-T cells, tumour cells, and healthy B cells over time, along with the corresponding dynamics of the equation terms related to competition, immunosuppression, and the effector-to-target ratio. The temporal dynamics of each CAR-T cell phenotype, along with the total target-antigen load, are presented in Figure SM.1.

**Figure 5.**
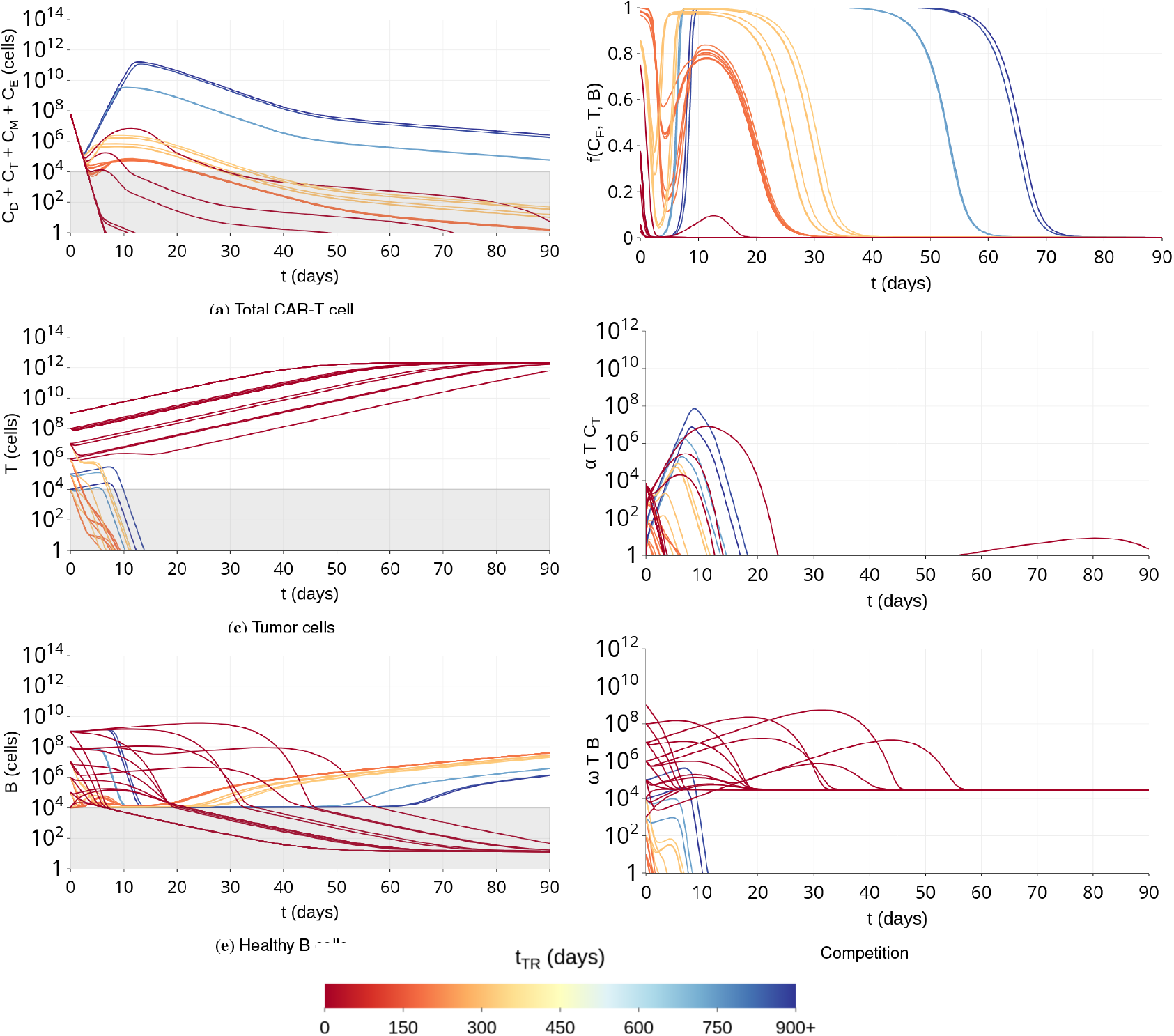
Temporal dynamics of total CAR-T cells (a), tumour cells (c), and healthy B cells (e) for *Profile CR90* under distinct theoretical relapse-free times (*t*_*TR*_). The impact of the initial target-antigen load composition is also reflected in the dynamic behaviour of specific biological mechanisms, including the effector-to-target ratio (b), immunosuppression (d), and competition (f).

The best therapeutic outcomes (longer relapse-free time – blueish curves) typically exhibit: (i) greater CAR-T cell expansion (Figure 5a), (ii) a rapid reduction of the target-antigen load below the clinical detection threshold within the first two weeks (Figure 5c and Figure 5e), (iii) prolonged BCA, followed by regrowth of healthy B cells, potentially restoring their homeostatic levels (Figure 5e), (iv) a more intense and sustained cytotoxic effect driven by the effector-to-target ratio, as described by the function *f* (*C*_*F*_, *T, B*) (Figure 5b), and (v) tumour-induced immunosuppression and competition between tumour and healthy B cells, particularly during the first two weeks after infusion (Figure 5d and Figure 5f). In contrast, when therapy fails immediately (*t*_*TR*_ = 0 – dark red curves), minimal cytotoxic effect is observed, with intense and persistent competition between cells expressing the target antigen, eventually allowing tumour cells to completely replace healthy B cells due to their competitive advantage.

Thus, therapy response is influenced by the initial composition of target-antigen-expressing cells, with optimal outcomes characterised by significant CAR-T cell expansion, rapid reduction of target antigens, prolonged BCA, and effective management of immunosuppression and competition, whereas therapy failure results from minimal cytotoxic effects and tumour cells replacing healthy B cells.

## Discussion

CAR-T cell immunotherapy has been highly successful in treating haematological cancers; however, high recurrence rates remain a significant challenge (1, 2). Further scientific research is needed to elucidate the various biological mechanisms involved in this therapy and to address unresolved issues (36). As a living drug, CAR-T cells exhibit distinct dynamics compared to chemotherapy drugs. Therefore, the description of pharmacokinetics and pharmacodynamics differs from traditional approaches, requiring the development of new methodologies (37). In addition to the complex multiphasic dynamics of CAR-T cells, the interaction of CAR-T cells with the patient’s immune system introduces an added layer of complexity. Healthy B cells, part of the immune system, also express the CD19 antigen targeted by CAR-T cells in many B cell malignancies. As new B cells are continuously generated in the bone marrow, this influx of antigen directly influences the dynamics of CAR-T cells, providing a continuous stimulus for expansion and cytotoxic activity, acting like an endogenous vaccine (38, 39). Interestingly, BCA represents a form of on-target, off-tumour toxicity that serves as a marker of successful therapy, as it indicates the presence of circulating functional CAR-T cells. Limited knowledge still exists regarding the impact of late BCA loss and its relationship with pre-infusion tumour burden in patients. Recently, Molinos-Quintana et al. (40) seek to clarify these issues by evaluating a group of 73 patients, reporting a positive correlation between BCA loss and CD19-positive relapse.

The importance of tumour burden in CAR-T cell immunotherapy outcomes is well documented in the literature (41), but the impact of healthy cells expressing the target antigen remains less understood. In this work, we developed a mathematical model to describe the interaction between various CAR-T cell phenotypes and target-antigen-expressing tumour and healthy B cells, with the aim of investigating how target-antigen-expressing cells influence the dynamics of immunotherapy. Healthy B cells can be subdivided into different stages of maturation, exhibiting variations in target antigen expression (42, 43). Due to the lack of experimental data on these subpopulations, and considering that the biological processes occurring in the marrow are reflected in the peripheral blood after a certain period, we chose to model a single population of healthy B cells. This encompasses all stages of B cell maturation that express the target antigen, and integrates the model alongside tumour cells and four CAR-T cell phenotypes, including functional (distributed and effector), memory, and exhausted CAR-T cells. Our model successfully captured distinct dynamics, such as varying phenotypic compositions within the total CAR-T cell population, with some patients experiencing prolonged BCA while others did not achieve it, ultimately resulting in diverse therapy responses. Despite the lack of experimental data for patients who achieved PR, SD, and PD responses, the generation of theoretical patient profiles demonstrated the model’s capability to describe these diverse response scenarios. The *κ*(*t*) function plays a fundamental role in incorporating heterogeneities from both the patient and the infused product into the dynamics. In summary, our model shows that patient and CAR-T cell product heterogeneities, the initial target-antigen load, and the corresponding proportions of tumour and healthy B cells are key factors that ultimately determine the outcome of the therapy.

As emphasised by Martínez-Rubio et al. (11), the attributes of CAR-T cell products are crucial in determining the outcomes of therapy. However, these are not the only factors at play; our findings suggest that they should also be considered alongside the composition of initial total target-antigen load. Unlike Martínez-Rubio et al., our results indicate that the dynamics of both CAR-T cells and tumour cells are influenced by the initial concentration of healthy B cells and the tumour burden expressing the target antigen.

A high number of healthy B cells may initially seem advantageous, as they promote the proliferation and reactivation of CAR-T cells without displaying immunosuppressive mechanisms. However, their impact is contingent upon the tumour burden at the time of infusion. An initial significantly high total number of target-antigen-expressing cells can adversely affect cytotoxic activity. Consequently, even if tumour burden initially decreases due to cytotoxicity, this reduction may be insufficient, allowing tumour cells to regrow and competitively exclude healthy B cells, potentially leading to tumour recurrence. As demonstrated by Molinos-Quintana et al. (40), a high tumour burden prior to infusion significantly influences therapeutic relapse.

We identified the critical role of the initial ratio between tumour and healthy target-antigen-expressing cells on the relapse-free time. Different ratios, including those with the same total antigen load, resulted in varied outcomes. In other words, adjustments to the baseline levels of tumour and healthy B cells were able to yield positive therapeutic results. Direct practical implications may arise from the fact that immunotherapy relapse is often preceded by the recovery of healthy B cells, indicating a loss or dysfunction of CAR-T cells. As evidenced by Molinos-Quintana et al. (40), patients with low pre-infusion tumor burden with loss of BCA have a potential treatment window. The combined conditions of a high concentration of healthy B cells and low tumour burden at relapse onset might create a favourable window for adjuvant therapeutic treatment as indicated by our results. Therefore, BCA monitoring is strongly recommended, as it can guide new treatments.

As noted by Brudno et al. (39) and Kochenderfer et al. (27), there is considerable heterogeneity in the dynamics among patients who achieve a complete response. Patient profiles vary in several aspects: the timing of the CAR-T cell peak (whether early or late), the expansion capacity of CAR-T cells (with differing peak concentrations), the dynamic behaviour of each CAR-T cell phase, and the duration of BCA. CAR-T cell immunotherapy is capable of inducing long-term remissions even after the loss of BCA, as shown by Kochenderfer et al. (27). Lee et al. (7) report that patients who achieved complete response did not experience prolonged BCA. Of the 115 patients reported by Wang et al. (9), two exhibited BCA lasting over a year, whereas the study published by Gardner et al. (5) reported an average BCA duration of 3 months for the patients. In Melenhorst et al.‘s milestone work, the two patients who have remained in remission for a decade achieved BCA, but with one of them showing increasing levels of healthy B cells. In addition to this diversity among responding patients, Fraietta et al. (45) highlight distinct temporal dynamic profiles between groups of patients with different responses to therapy. This justifies the analyses performed in this study to observe the impact of healthy B cells on the therapy outcome. As highlighted by Kröger et al. (6), monitoring B-cell aplasia alongside the status of remission and the persistence of CAR-T cells can guide clinical decisions regarding disease evolution and the need for additional treatments. Our model assisted in exploring the role of the phenotypic composition of the total CAR-T cell population, which is predominantly composed of effector CAR-T cells at peak time, but displays varied behaviour in terms of memory and exhausted CAR-T cell formation. As demonstrated, our model has a strong capacity to describe different dynamic profiles. In the various evaluated scenarios, when describing the temporal dynamics of healthy B cells, we highlighted the wide range of values assumed by the parameters that describe the growth rate of healthy B cells and the influx of new cells from the marrow. This appears clinically plausible, as the heterogeneity among patients can be reflected in these parameters. It is important to note that patients undergoing CAR-T cell immunotherapy have typically received multiple lines of treatment, some of them presenting refractory-relapsed cancers, which may affect the characteristics of their B cell populations. Target antigen is a key factor in CAR-T cell immunotherapy. Here, we evaluated CAR T cells targeting the B cell protein CD19; however, we emphasise that the application of our model can be extended to cover other target antigens, such as CD22, and dual CARs, a strategy currently well-utilised in cases of resistance (46).

In this work, we were able to evaluate various aspects associated with BCA and the cells expressing the target antigen in response to therapy, thanks to the integration of healthy B cells into the mathematical modelling. While mathematical and computational modelling proved useful in these analyses, it is important to mention that it is not a common practice in clinical medicine, as reported in (47). The lack of openly shared clinical data is one of the major challenges in CAR-T cell immunotherapy modelling (48). We encountered significant difficulties in acquiring patient data to calibrate the developed model. Most available studies do not provide the temporal dynamics of healthy B cells, merely reporting the duration of BCA. Moreover, data may be reported in different units for total CAR-T cells, tumour cells, and healthy B cells. Various experimental factors affect these measurements, and there is no standardised method in the literature to convert these units into absolute cell counts. Despite these limitations, we developed a model capable of describing some aspects of the heterogeneity present in CAR-T cell immunotherapy and generating insights that can contribute to a better understanding of the importance of BCA and the influence of target-antigen-expressing tumour and healthy B cells.

## ACKNOWLEDGEMENTS

Emanuelle A. Paixão is supported by a post-doctoral fellowship from the Institutional Training Program (PCI) at LNCC of the Brazilian National Council for Scientific and Technological Development (CNPq), Grant 301998/2023-0. Regina C. Almeida acknowledges the support provided by CNPq, Grant 306588/2022-6. Artur C. Fassoni was supported by Alexander von Humboldt Foundation and Coordenação de Aperfeiçoamento de Pessoal de Nível Superior - Brasil (CAPES) - Finance Code 001, and partially supported by FAPEMIG RED-00133-21.

