## Supplementary material for "Healthy B cells: allies or adversaries of CAR-T cell immunotherapy?": supplementary_material.pdf

**This Supplementary Material (SM) is organized as follows. Section S.1 describes additional characteristics of the developed model. Section S.2 presents patient data collected from studies (1, 2). Section S.3 provides complementary results and model simulations for each patient from Kochenderfer et al. (2). Finally, we present the calibrated parameter values used to simulate.**

### Mathematical Model

Tumour and healthy B cell populations grow subject to both intraspecific and interspecific competition. The former is modelled by a logistic growth term, which indicates limitations in available resources within the tumour microenvironment. Both populations grow logistically, each with its respective intrinsic growth rate ($r_1$ for tumour cells and $r_2$ for healthy B cells), and they share the same carrying capacity ($1/b$). Interspecific competition captures the effect of one population on the other. Thus, the dynamics of the tumour cell population is given by

$$\frac{\mathrm{d}T}{\mathrm{d}t} = r_1 T\left[1 - b(T + g(B))\right],$$

in which the growth term is defined with a maximum growth rate of $r_1$, a carrying capacity of $1/b$, and the function $g(B)$ models the impact of healthy B cells on tumour cell growth. By introducing the competition coefficient $w_1$, which incorporates the carrying capacity, we can rewrite the previous equation as

$$\frac{\mathrm{d}T}{\mathrm{d}t} = r_1 T(1 - bT) - r_1 \omega_1 T B\,.$$

Considering that tumour cells have a stronger competitive advantage over healthy B cells (3), we assumed $\omega_1 \approx 0$, which leads to

$$\frac{\mathrm{d}T}{\mathrm{d}t} = r_1 T(1 - bT)\,.$$

In the same line, we assume that tumour cells have a negative impact on the growth of healthy B cells. Defining the competition coefficient $\omega$, the dynamics of healthy B cells is given by

$$\frac{\mathrm{d}B}{\mathrm{d}t} = r_2 B(1 - bB) - \omega T B\,.$$

### Experimental data

Tables 1–3 present patient data collected from Lee et al. (1) and Kochenderfer et al. (2).

Data from Lee et al. encompass median values from a group of responding patients, including time points for the abundance of total CAR-T cells, tumour cells, and healthy B cells in peripheral blood. The data from Kochenderfer et al. are from patients with different subtypes of diffuse large B-cell lymphoma (DLBCL) who achieved complete response. These data include time points for total CAR-T cell and healthy B cells. CAR-T cell data are available for up to 90 days post-infusion, depending on the patient. In contrast, data for healthy B cells are concentrated at later stages in the dynamics, although baseline values prior to therapy are available for some patients.

### Simulation Results

Here, we present additional results for *Profile CR90*, the median patient from (1). Figure 1 illustrates the temporal dynamics of functional, memory, and exhausted CAR-T cells, along with the total target-antigen load, for various combinations of initial conditions ($T(0)$ and $B(0)$). These combinations lead to different relapse-free times. Therapies with better outcomes tend to exhibit higher peaks of functional CAR-T cells (Figure 1a), and increased exhaustion and memory cell generation (Figures 1b and 1c). Higher levels of total target-antigen load are observed in non-responder patients (Figure 1d).

We now present model simulations for four patients reported in (2). During calibration, we observed that parameters associated with healthy B cells ($\gamma_2$, $r_2$, and $B_p$) significantly influence the long-term dynamics. Adjusting these parameters enabled control over the duration and loss of BCA. The analysed patients exhibit the following key characteristics:

- high initial CAR-T cell expansion rates ($1.97 \leq r_{min} + p_1 \leq 3.65$) and baseline expansion levels ($1.0 \times 10^{-2} \leq r_{min} \leq 6.0 \times 10^{-1}$);
- at its peak value, the total CAR-T cell population is primarily composed of effector CAR-T cells;
- the formation of a memory pool clinically detectable in 3 out of 4 patients;

**Table 1.** Data on the abundance of total CAR-T cells, tumour cells, and healthy B cells in peripheral blood extracted from Lee et al. (1), using the software WebPlotDigitizer (4). These are median values for a group of responding patients, with the number of patients indicated in parentheses.

| Days after infusion | Total CAR-T cell counts | Tumour cells | Healthy B cells |
|---|---|---|---|
| Responding patient group | | | |
| -1 | – | 1,367,545.0 (14) | – |
| 0 | – | – | 22,421,321.0 (14) |
| 3 (1-5) | 134,898.0 (14) | 320,871.0 (14) | 5,345,621.0 (14) |
| 7 (5-9) | 19,898,391.0 (14) | 10,000.0* (14) | 1,561,602.0 (14) |
| 14 (12-16) | 20,128,344.0 (14) | 10,000.0* (11) | 10,000.0* (12) |
| 28 (25-31) | 1,229,228.0 (14) | 10,000.0* (11) | 10,000.0*# (12) |
| 47 (45-49) | 10,000.0* (6) | – | 54,676,675.0# (6) |
| 68 (55-81) | 10,000.0* (2) | – | 1,825,097.0# (3) |

*Data below the detection threshold (assumed equal to $10^4$ cells).
#Existence of patients with normal B-cell progenitors presence.

**Table 2.** CAR-T cell data extracted from Kochenderfer et al. (2), whose patients achieved complete response, with the day of the last follow-up measurement indicated in parentheses. The data were obtained using the software WebPlotDigitizer (4).

| Days after infusion | CAR-T cells/$\mu$L | Total CAR-T cell counts |
|---|---|---|
| Patient K2 (PMBCL) — CAR-T dose = $6.0\times10^8$ cells — CR(1680) | | |
| 5 | 3.1776 | $1.5888\times10^9$ |
| 8.1915 | 59.2523 | $2.9626\times10^{10}$ |
| 12.2340 | 5.9813 | $2.9907\times10^9$ |
| 14.1489 | 7.1028 | $3.5514\times10^9$ |
| 32.8723 | 1.4953 | $7.4766\times10^8$ |
| Patient K7 (DLBCL NOS) — CAR-T dose = $1.50\times10^8$ cells cells — CR(1530) | | |
| 10 | 3.8189 | $1.9094\times10^9$ |
| 12.0455 | 6.8802 | $3.4401\times10^9$ |
| 14.0909 | 8.1233 | $4.0616\times10^9$ |
| 17.0455 | 8.1680 | $4.0840\times10^9$ |
| 40 | 3.0613 | $1.5306\times10^9$ |
| 68.4091 | 3.4917 | $1.7459\times10^9$ |
| Patient K8 (PMBCL) — CAR-T dose = $1.50\times10^8$ cells — CR(1140) | | |
| 5.9091 | 9.6970 | $4.8485\times10^9$ |
| 8.8636 | 15.7576 | $7.8788\times10^9$ |
| 13.1818 | 2.4242 | $1.2121\times10^9$ |
| 15.6818 | 1.8182 | $9.0909\times10^8$ |
| Patient K15 (DLBCL NOS) — CAR-T dose = $6.0\times10^7$ cells — CR(1260) | | |
| 5.6857 | 73.4000 | $3.6700\times10^{10}$ |
| 8.0769 | 765.7143 | $3.8286\times10^{11}$ |
| 15.0571 | 97.4000 | $4.8700\times10^{10}$ |
| 26.0286 | 61.4000 | $3.0700\times10^{10}$ |
| 57.3429 | 11.0000 | $5.5000\times10^9$ |
| 87.9714 | 2.6000 | $1.3000\times10^9$ |

DLBCL — diffuse large B-cell lymphoma; NOS — not otherwise specified; PMBCL — primary mediastinal B cell lymphoma.

- occurrence of BCA with variable duration among patients.

The healthy B cell population reaches a temporary equilibrium before the CAR-T cell peak. A few days after therapy, CAR-T cells eliminate healthy B cells, resulting in BCA. However, after a period that varies from patient to patient, CAR-T cells can no longer prevent healthy B cells from resuming growth. As these cells expand, they eventually become detectable.

The analysed responding patients exhibited high peaks of CAR-T cells, composed primarily of the effector phenotype, along with a persistent memory cell population. All patients experienced BCA driven by CAR-T cell therapy, with only one maintained BCA until the final day of analysis. The tumour burden remained low, below the detection threshold, until the final day of follow-up, preventing the identification of any competitive advantage of the tumour over healthy B cells.

Patient K2 exhibits the smoothest distribution phase

**Table 3.** Healthy B cell data extracted from Kochenderfer et al. (2), whose patients achieved complete response, with the day of the last follow-up measurement indicated in parentheses. The data were obtained using the software WebPlotDigitizer (4).

| **Days after infusion** | **Healthy B cells/$\mu$L** | **Healthy B cell counts** |
|---|---|---|
| Patient K2 (PMBCL) — CR(1680) | | |
| 327.9503 | 17.9389 | $8.9695 \times 10^{9}$ |
| 389.4410 | 21.3740 | $1.0687 \times 10^{10}$ |
| 450.9317 | 28.2443 | $1.4122 \times 10^{10}$ |
| 568.3230 | 30.5344 | $1.5267 \times 10^{10}$ |
| 691.3043 | 48.8550 | $2.4427 \times 10^{10}$ |
| 808.6957 | 63.7405 | $3.1870 \times 10^{10}$ |
| 870.1863 | 60.3053 | $3.0153 \times 10^{10}$ |
| 959.6273 | 63.7405 | $3.1870 \times 10^{10}$ |
| 1110.5590 | 80.9160 | $4.0458 \times 10^{10}$ |
| 1322.9814 | 66.0305 | $3.3015 \times 10^{10}$ |
| 1501.8634 | 83.2061 | $4.1603 \times 10^{10}$ |
| 1714.2857 | 53.4351 | $2.6718 \times 10^{10}$ |
| Patient K7 (DLBCL NOS) — CR(1530) | | |
| 272.2222 | 29.5455 | $1.4773 \times 10^{10}$ |
| 333.3333 | 39.7727 | $1.9886 \times 10^{10}$ |
| 422.2222 | 78.4091 | $3.9205 \times 10^{10}$ |
| 538.8889 | 121.5909 | $6.0795 \times 10^{10}$ |
| 777.7778 | 152.2727 | $7.6136 \times 10^{10}$ |
| 838.8889 | 181.8182 | $9.0909 \times 10^{10}$ |
| 927.7778 | 189.7727 | $9.4886 \times 10^{10}$ |
| 1022.2222 | 211.3636 | $1.0568 \times 10^{11}$ |
| 1172.2222 | 203.4091 | $1.0170 \times 10^{11}$ |
| 1350 | 195.4545 | $9.7727 \times 10^{10}$ |
| 1555.5556 | 159.0909 | $7.9545 \times 10^{10}$ |
| Patient K8 (PMBCL) — CR(1140) | | |
| 0 | 21.5909 | $1.0795 \times 10^{10}$ |
| 272.0497 | 1.1364 | $5.6818 \times 10^{8}$ |
| 389.441 | 0.0 | 0.0 |
| 657.764 | 0.0 | 0.0 |
| 1021.1180 | 0.0 | 0.0 |
| 1200 | 0.0 | 0.0 |
| Patient K15 (DLBCL NOS) — CR(1260) | | |
| 0 | 146.1832 | $7.3092 \times 10^{10}$ |
| 26.0870 | 1.9084 | $9.5420 \times 10^{8}$ |
| 54.0373 | 1.9084 | $9.5420 \times 10^{8}$ |
| 87.5776 | 1.9084 | $9.5420 \times 10^{8}$ |
| 450.9317 | 102.6718 | $5.1336 \times 10^{10}$ |
| 540.3727 | 166.7939 | $8.3397 \times 10^{10}$ |
| 657.7640 | 161.0687 | $8.0534 \times 10^{10}$ |
| 836.6460 | 185.1145 | $9.2557 \times 10^{10}$ |
| 959.6273 | 151.9084 | $7.5954 \times 10^{10}$ |
| 959.6273 | 138.1679 | $6.9084 \times 10^{10}$ |
| 959.6273 | 193.1298 | $9.6565 \times 10^{10}$ |

DLBCL — diffuse large B-cell lymphoma; NOS — not otherwise specified; PMBCL — primary mediastinal B cell lymphoma.

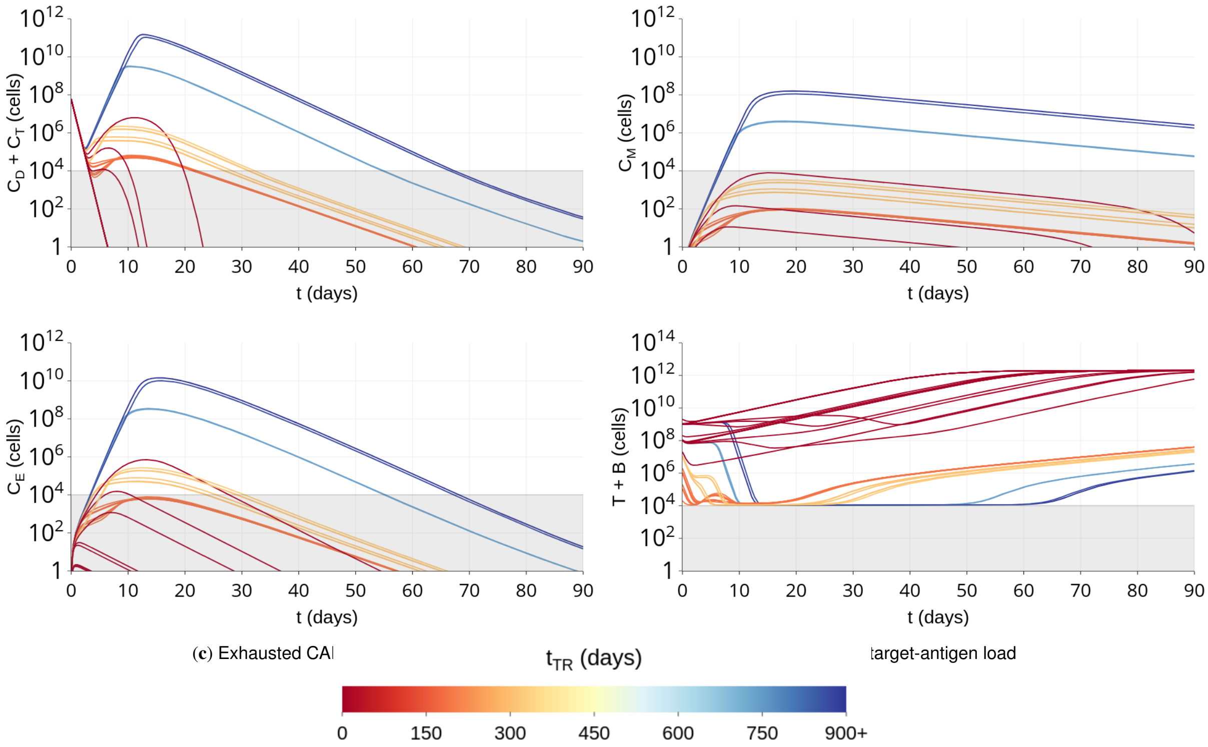


**Figure 1.** Temporal dynamics of functional CAR-T cells (a), memory CAR-T cells (b), exhausted CAR-T cells (c), and the total target-antigen load for *Profile CR90* under distinct theoretical relapse-free times ($t_{TR}$).

with $5.1194 \times 10^7$ engrafted cells (Figure 2). In the early stages of the dynamics, we observed a sharp decline in populations expressing the target antigen. Following this decline, tumour cells remain at very small values within the undetectable range, indicating a complete response. During the distribution phase, a slight increase is observed in the healthy B cell population; however, this growth is halted by the engrafted CAR-T cells, bringing the population into a period of temporary equilibrium. BCA is sustained until day 919. The $\kappa(t)$ function displays a plateau, followed by a rapid decline to the minimum basal expansion level. The length of this plateau closely corresponds to the timing of the distribution and expansion phases in the overall total CAR-T cell population.

The simulation results for Patient K7 are presented in Figure 3. As shown, CAR-T cells were unable to control the growth of healthy B cells during the early stages of the dynamics. Compared to the dynamics of patient K2, patient K7 shows: (i) a smoother transition between the expansion and contraction phases of CAR-T cells, with a later peak; (ii) minimal exhaustion, as the memory cell subpopulation consistently exceeds that of exhausted cells; and (iii) a distinct profile in the $\kappa(t)$ function, lacking a plateau and showing a slow decline to the minimum basal expansion level. This contrasts with patient K2, where the total CAR-T cell population peaked earlier.

The similarity between the peaks of the memory and exhausted CAR-T cell subpopulations in the model dynamics for patient K8 is remarkable (Figure 4. However, memory CAR-T cells exhibit a significantly lower mortality rate, allowing them to be detected during the persistence phase. Notably, patient K8 was the only one among the four to sustain BCA throughout the analysis period, lasting over 3 years. The CAR-T cell expansion was the largest among all patients analysed, as indicated in the $\kappa(t)$ profile shown in Figure 4c.

Finally, the model dynamics for patient K15 are shown in Figure 5. Total CAR-T cells exhibit the highest peak but remain undetectable during the persistence phase with memory cells peaking in the non-detection region. The profile of the $\kappa(t)$ function resembles that of patient K8 but features a longer duration of maximum expansion, a faster decay, and a higher baseline minimum level. The loss of BCA occurs after approximately a year (356 days). Among the four patients, the tumour cell population exhibits a slower initial decline in its dynamics.

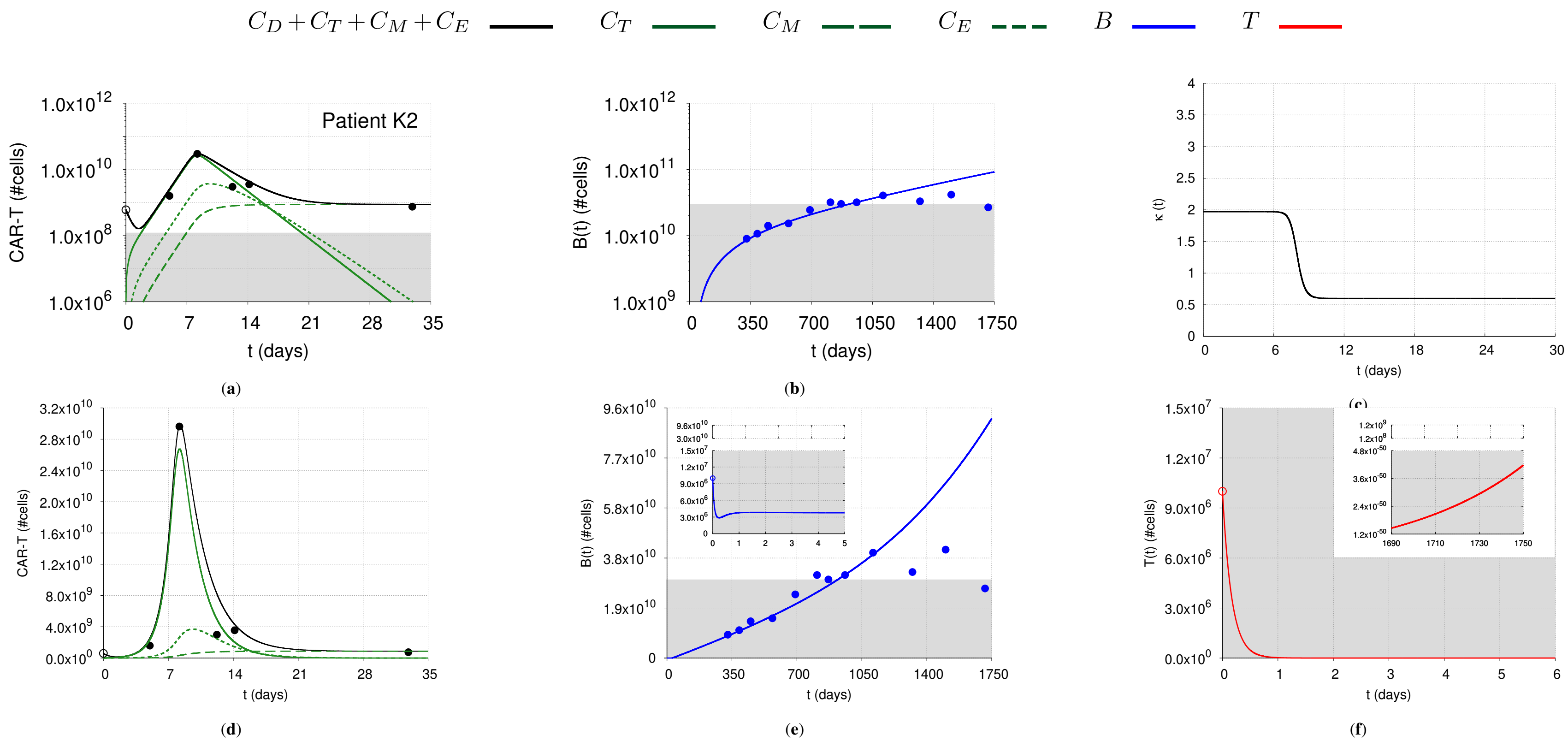


**Figure 2.** Model simulation for DLBCL Patient K2 reported in Kochenderfer et al. (2), whose data are shown by circular dots and estimated initial values are indicated by empty circles. The total CAR-T cell population (▬) is divided into effector ($C_T$), memory ($C_M$), and exhausted ($C_E$) phenotypes, shown in continuous, dashed, and dotted green lines, respectively. Tumour cells (▬) decay exponentially due to the cytotoxic effect of CAR-T cells but show a growth trend by day 1,680. Healthy B cells (▬) remain undetectable until the loss of BCA, which occurs after two years. Gray shadows represent undetectable levels ($\leq$ 1.1850 $\times 10^8$ cells for CAR-T cells and 3.0 $\times 10^{10}$ cells for healthy B cells). The temporal dynamics of each population are presented in logarithmic (top panel) and linear (bottom panel) scales. The corresponding time-dependent expansion rate function ($\kappa(t)$) is shown in (**c**).

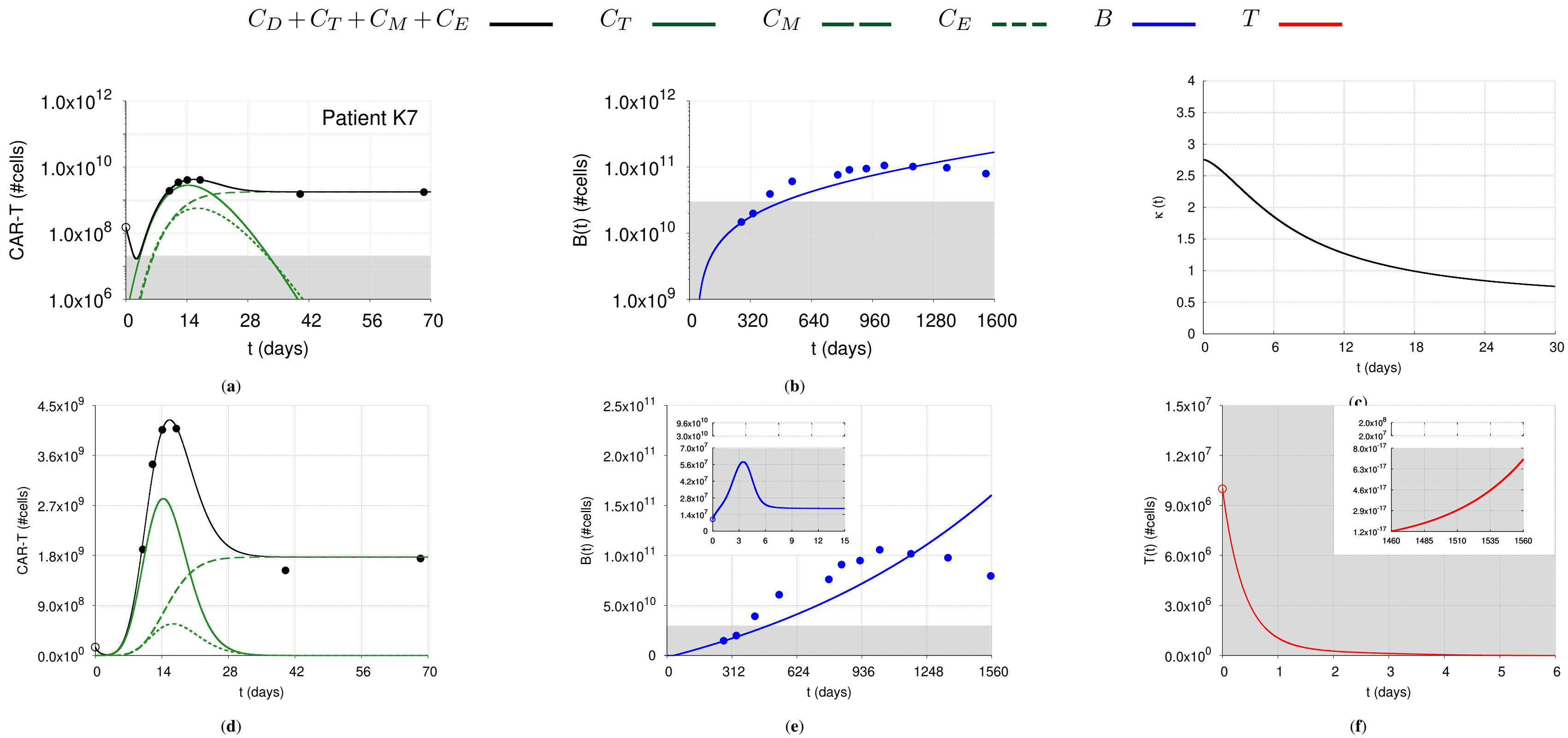


**Figure 3.** Model simulation for Patient K7 reported in Kochenderfer et al. (2), whose data are shown by circular dots and estimated initial values are indicated by empty circles. The total CAR-T cell population (—) is divided into effector ($C_T$), memory ($C_M$), and exhausted ($C_E$) phenotypes, shown in continuous, dashed, and dotted green lines, respectively. Tumour cells (—) decay exponentially due to the cytotoxic effect of CAR-T cells but show a growth trend by day 1,530. Healthy B cells (—) remain undetectable until the loss of BCA which occurs after one year. Gray shadows represent undetectable levels ($\leq 2.0420 \times 10^7$ cells for CAR-T cells and $3.0 \times 10^{10}$ cells for healthy B cells). The temporal dynamics of each population are presented in logarithmic (top panel) and linear (bottom panel) scales. The corresponding time-dependent expansion rate function ($\kappa(t)$) is shown in (**c**).

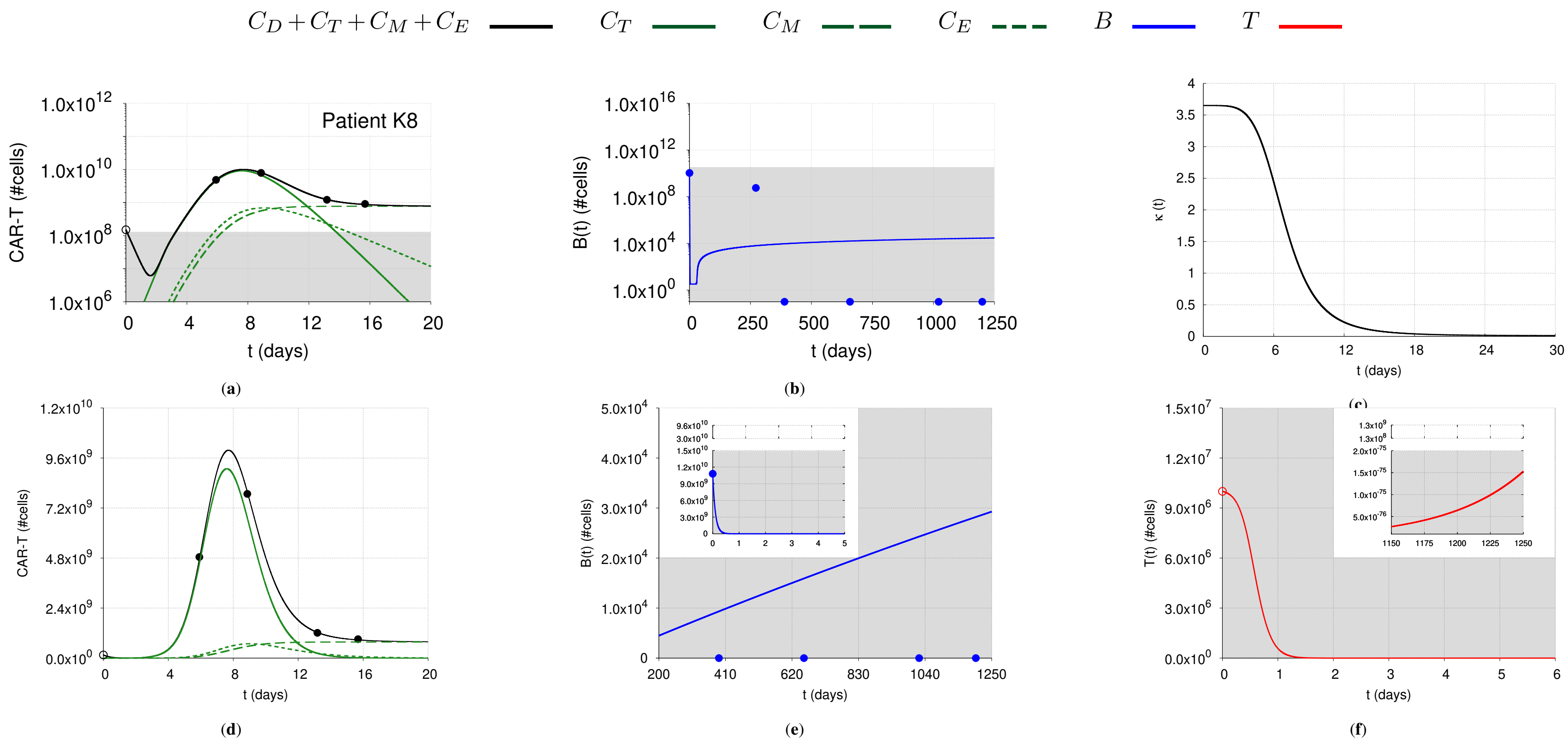


**Figure 4.** Model simulation for Patient K8 reported in Kochenderfer et al. (2), whose data are shown by circular dots and estimated initial values are indicated by empty circles. The total CAR-T cell population (▬) is divided into effector ($C_T$), memory ($C_M$), and exhausted ($C_E$) phenotypes, shown in continuous, dashed, and dotted green lines, respectively. Tumour cells (▬) decay due to the cytotoxic effect of CAR-T cells but show a growth trend by day 1,140. Healthy B cells (▬) remain undetectable throughout the simulation. Gray shadows represent undetectable levels ($\leq$ 1.3394 $\times 10^8$ cells for CAR-T cells and 3.0 $\times 10^{10}$ cells for healthy B cells). The temporal dynamics of each population are presented in logarithmic (top panel) and linear (bottom panel) scales. The corresponding time-dependent expansion rate function ($\kappa(t)$) is shown in (**c**).

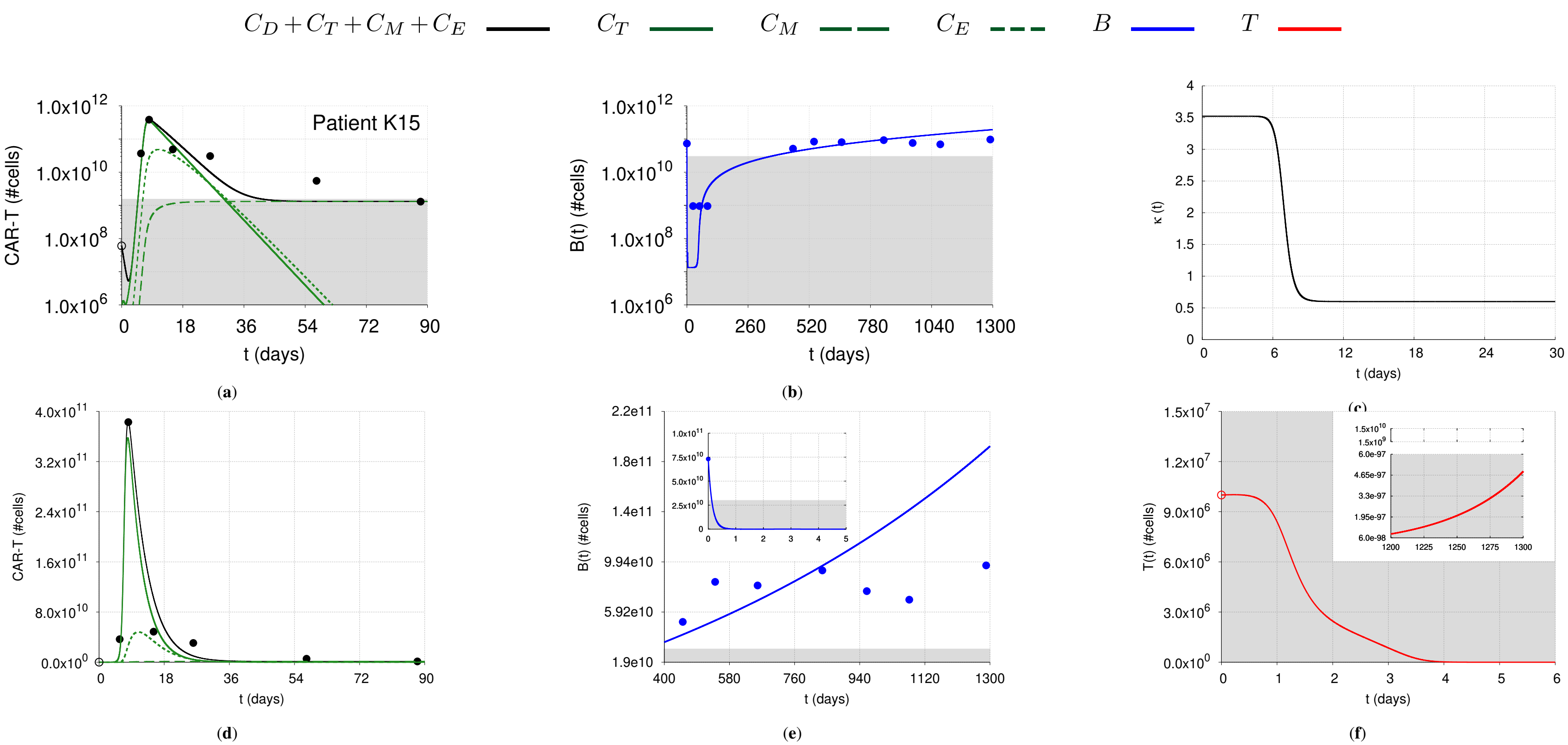


**Figure 5.** Model simulation for Patient K15 reported in Kochenderfer et al. (2), whose data are shown by circular dots and estimated initial values are indicated by empty circles. The total CAR-T cell population (▬) is divided into effector ($C_T$), memory ($C_M$), and exhausted ($C_E$) phenotypes, shown in continuous, dashed, and dotted green lines, respectively. Tumour cells (▬) reach very low levels from day 4 onwards but show a growth trend around day 1,260. Healthy B cells (▬) remain undetectable until the loss of BCA, which occurs after one year. Gray shadows represent undetectable levels ($\leq$ 1.5314 $\times 10^9$ cells for CAR-T cells and 3.0 $\times 10^{10}$ cells for healthy B cells). The temporal dynamics of each population are presented in logarithmic (top panel) and linear (bottom panel) scales. The corresponding time-dependent expansion rate function ($\kappa(t)$) is shown in (**c**).

**Table 4.** Calibrated parameter values, in appropriate units (a.u.), used in the simulations for responding group (1) (*Profile CR90*), theoretical patients (*Profiles PR90* and *PD90*), and real patients (2) (*Patients K2*, *K7*, *K8*, and *K15* ). The initial conditions are specified, with the effector, memory, and exhausted CAR-T phenotypes set to zero, as phenotypic differentiation occurs throughout the evolution of the dynamics. The approximated total number of engrafted CAR-T cells ($EC$) is evaluated for each patient using $EC \approx \left(\frac{\eta}{\beta+\eta}\right) C_D(0)$. Parameters whose values were the same for all patients are: $\alpha = 5.5 \times 10^{-7}$, $1/b = 2.0 \times 10^{12}$, and $\vartheta = 1.0$.

| Parameter | Profile CR90 | Profile PR90 | Profile PD90 | Patient K2 | Patient K7 | Patient K8 | Patient K15 |
|---|---|---|---|---|---|---|---|
| $\beta$ | 2.8 | 2.8 | 2.8 | 1.39363 | 1.19363 | 2.39363 | 1.39363 |
| $\eta$ | $1.2 \times 10^{-4}$ | $1.2 \times 10^{-4}$ | $1.2 \times 10^{-4}$ | $1.3 \times 10^{-1}$ | $1.3 \times 10^{-2}$ | $1.0 \times 10^{-2}$ | $1.3 \times 10^{-1}$ |
| $r_{min}$ | $1.0 \times 10^{-1}$ | $1.0 \times 10^{-3}$ | $1.0 \times 10^{-3}$ | $6.0 \times 10^{-1}$ | $4.87 \times 10^{-1}$ | $1.0 \times 10^{-2}$ | $6.0 \times 10^{-1}$ |
| $p_1$ | 1.91 | 1.2015 | 1.0822 | 1.37 | 2.27 | 3.64 | 2.922 |
| $p_2$ | $7.1429 \times 10^{-2}$ | $8.3333 \times 10^{-2}$ | $5.0 \times 10^{-2}$ | $1.25 \times 10^{-1}$ | $1.2658 \times 10^{-1}$ | $1.4493 \times 10^{-1}$ | $1.4388 \times 10^{-1}$ |
| $p_3$ | 5.5 | 5.5 | $1.8 \times 10^{1}$ | $2.5 \times 10^{1}$ | 1.52 | 5.0 | $1.8 \times 10^{1}$ |
| $A$ | $5.0 \times 10^{4}$ | $5.0 \times 10^{4}$ | $5.0 \times 10^{4}$ | $1.0 \times 10^{1}$ | $1.0 \times 10^{1}$ | 1.0 | $1.0 \times 10^{1}$ |
| $\xi$ | $2.9 \times 10^{-1}$ | $2.9 \times 10^{-1}$ | $2.9 \times 10^{-1}$ | $9.59828 \times 10^{-1}$ | $9.87828 \times 10^{-1}$ | $9.99828 \times 10^{-1}$ | $7.79828 \times 10^{-1}$ |
| $\epsilon$ | $2.5 \times 10^{-4}$ | $2.5 \times 10^{-4}$ | $2.5 \times 10^{-4}$ | $8.0 \times 10^{-3}$ | $5.5 \times 10^{-2}$ | $2.1 \times 10^{-2}$ | $6.5 \times 10^{-4}$ |
| $\lambda$ | $5.0 \times 10^{-2}$ | $5.0 \times 10^{-2}$ | $5.0 \times 10^{-2}$ | $1.0 \times 10^{-1}$ | $1.0 \times 10^{-1}$ | $5.0 \times 10^{-2}$ | $7.0 \times 10^{-2}$ |
| $\theta$ | $7.0 \times 10^{-13}$ | $7.0 \times 10^{-13}$ | $7.0 \times 10^{-13}$ | $5.0 \times 10^{-14}$ | $5.0 \times 10^{-15}$ | $5.0 \times 10^{-14}$ | $5.0 \times 10^{-15}$ |
| $\mu$ | $6.2 \times 10^{-2}$ | $6.2 \times 10^{-2}$ | $6.2 \times 10^{-2}$ | $4.0 \times 10^{-6}$ | $1.0 \times 10^{-6}$ | $1.0 \times 10^{-5}$ | $4.0 \times 10^{-6}$ |
| $\delta$ | $3.5 \times 10^{-1}$ | $3.5 \times 10^{-1}$ | $3.5 \times 10^{-1}$ | $4.5 \times 10^{-1}$ | $4.5 \times 10^{-1}$ | $4.5 \times 10^{-1}$ | $2.8 \times 10^{-1}$ |
| $\gamma_1$ | 2.5 | 2.5 | 2.5 | 6.19 | 2.85 | 7.19 | 6.09 |
| $\gamma_2$ | 2.6 | 2.6 | 2.6 | 7.2 | 2.9 | 8.2 | 6.5 |
| $r_1$ | $1.76 \times 10^{-1}$ | $1.76 \times 10^{-1}$ | $1.76 \times 10^{-1}$ | $1.76 \times 10^{-2}$ | $1.76 \times 10^{-2}$ | $1.76 \times 10^{-2}$ | $1.76 \times 10^{-2}$ |
| $r_2$ | $6.98 \times 10^{-2}$ | $6.98 \times 10^{-2}$ | $6.98 \times 10^{-2}$ | $1.0 \times 10^{-3}$ | $8.2 \times 10^{-4}$ | $1.0 \times 10^{-4}$ | $1.0 \times 10^{-3}$ |
| $B_p$ | $2.7 \times 10^{4}$ | $2.7 \times 10^{4}$ | $2.7 \times 10^{4}$ | $2.7 \times 10^{7}$ | $5.5 \times 10^{7}$ | $2.7 \times 10^{1}$ | $8.7 \times 10^{7}$ |
| $\omega$ | $1.0 \times 10^{-9}$ | $1.0 \times 10^{-9}$ | $1.0 \times 10^{-9}$ | $1.0 \times 10^{-6}$ | $1.0 \times 10^{-7}$ | $1.0 \times 10^{-6}$ | $7.0 \times 10^{-7}$ |
| Initial Condition | | | | | | | |
| $C_D(0)$ (total dose) | $6.0 \times 10^{7}$ | $6.0 \times 10^{7}$ | $6.0 \times 10^{7}$ | $6.0 \times 10^{8}$ | $1.5 \times 10^{8}$ | $1.5 \times 10^{8}$ | $6.0 \times 10^{7}$ |
| $T(0)$ | $1.36754487 \times 10^{6}$ | $1.36754487 \times 10^{6}$ | $1.36754487 \times 10^{6}$ | $1.0 \times 10^{7}$ | $1.0 \times 10^{7}$ | $1.0 \times 10^{7}$ | $1.0 \times 10^{7}$ |
| $B(0)$ | $2.2421320913 \times 10^{7}$ | $2.2421320913 \times 10^{7}$ | $2.2421320913 \times 10^{7}$ | $1.0 \times 10^{7}$ | $1.0 \times 10^{7}$ | $1.0795 \times 10^{10}$ | $7.3092 \times 10^{10}$ |
| Engrafted CAR-T Cells | | | | | | | |
| $EC$ | $2.5713 \times 10^{3}$ | $2.5713 \times 10^{3}$ | $2.5713 \times 10^{3}$ | $5.1193531 \times 10^{7}$ | $1.61607 \times 10^{5}$ | $6.24056 \times 10^{5}$ | $5.119353 \times 10^{6}$ |
